# Mechanistic Insights into the Active Site and Allosteric Communication Pathways in Human Nonmuscle Myosin-2C

**DOI:** 10.1101/138891

**Authors:** Krishna Chinthalapudi, Sarah M. Heissler, Matthias Preller, James R. Sellers, Dietmar J. Manstein

## Abstract

The cyclical interaction of myosin with F-actin and nucleotides is the basis for contractility of the actin cytoskeleton. Despite a generic, highly conserved motor domain, ATP turnover kinetics and their activation by F-actin vary greatly between myosins-2 isoforms. Here, we present a 2.25 Å crystal structure of the human nonmuscle myosin-2C motor domain, one of the slowest myosins characterized. In combination with integrated mutagenesis, ensemble-solution kinetics, and molecular dynamics simulations approaches, this study reveals an allosteric communication pathway that connects the distal end of the motor domain with the active site. Genetic disruption of this pathways reduces nucleotide binding and release kinetics up to 85-fold and abolishes nonmuscle myosin-2 specific kinetic signatures. These results provide insights into structural changes in the myosin motor domain that are triggered upon F-actin binding and contribute critically to the mechanochemical behavior of stress fibers, actin arcs, and cortical actin-based structures.

## Introduction

The extent to which filamentous actin (F-actin) can activate the enzymatic output of conventional myosins-2 varies by more than two orders of magnitude (1, 2). Despite a substantial sequence identity and a highly conserved actomyosin ATPase cycle, structural and allosteric adaptations causative for the tremendous enzymatic and hence physiological differences are largely unknown (**Figure 1A**) (1, 3).

**Figure 1:**
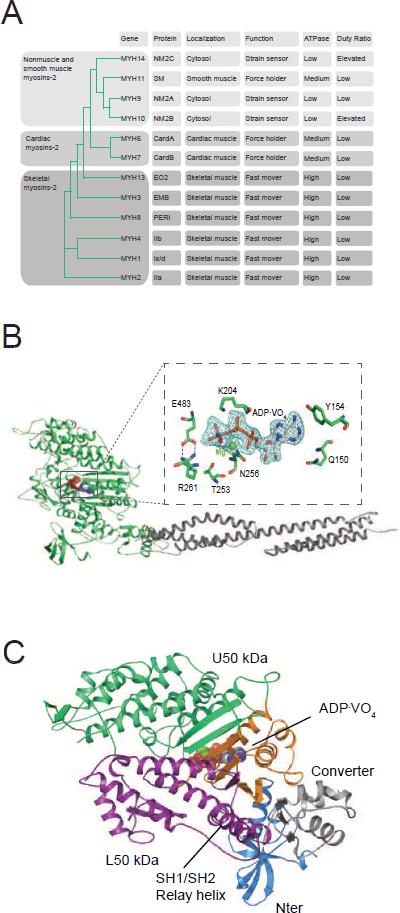
Myosin-2 phylogeny, overall topology, and active site characteristics of human NM2C. (**A**) Phylogenetic analysis divides human myosins-2 in the three subfamilies (i) nonmuscle and smooth muscle myosins-2, (ii) cardiac, (iii) and skeletal muscle myosins-2 (52). Nonmuscle myosin-2s are essential for the structural integrity of the cytoplasmic architecture during cell shape remodeling and motile events of eukaryotic cells whereas all other myosins-2 play eminent roles in the contraction of smooth, cardiac and striated muscle cells (53). Abbreviations used: NM2A: nonmuscle myosin-2A, NM2B: nonmuscle myosin-2B; NM2C: nonmuscle myosin-2C; SM: smooth muscle myosin-2; CardA: α-cardiac myosin-2; CardB: β-cardiac myosin-2; EO2: extraocular myosin-2; EMB: embryonic myosin-2; PERI: perinatal myosin-2; IIb: fast skeletal muscle myosin-2; IIx/d: skeletal muscle myosin-2; IIa: slow skeletal muscle myosin-2. (**B**) Architecture of the crystallized NM2C construct in the pre-powerstroke state. The myosin motor domain and the α-actinin repeats are shown in cartoon representation in green and grey color. The nucleotide is shown in spheres representation. *Inset*, Conserved key residues that interact with the nucleotide in the NM2C active site. The F_o_-F_c_ omit map of Mg^2+^ ·ADP·VO_4_ is contoured at 4σ. The salt bridge between switch-1 R261 and switch-2 E483 is highlighted. (**C**) Subdomain architecture of NM2C. The U50 kDa is shown in green, the L50 kDa in purple, the converter in grey, and the Nter in blue. The region shown in orange corresponds to the active site and the junction of U50 kDa and L50 kDa. The bound nucleotide is shown in spheres representation. The location of the SH1-SH2 helix and the relay helix in the L50 kDa is highlighted.

Nonmuscle myosin-2C is one of the slowest myosins-2 as its steady-state ATPase activity lacks potent F-actin activation (4). Transient kinetic signatures of the nonmuscle myosin-2C enzymatic cycle include a high affinity for ADP, small kinetic (**k_−AD_/k**_−D_) and thermodynamic (**K_AD_/K**_D_) coupling ratios, and a higher duty ratio than it is commonly found in myosins-2 (**Figure 1A - Figure supplement 1A**). Together, these signatures qualify nonmuscle myosin-2C as a dynamic strain-sensing actin tether (3-5). The participation in the active regulation of cytoplasmic contractility in cellular processes including cytokinesis, neuronal dynamics, adhesion, and tension maintenance are in line with this interpretation (6-8). Nonmuscle myosin-2C has recently received great attention as the near atomic resolution cryo electron microscopic structure of its motor domain bound to F-actin has been determined (9). The structure of the actomyosin complex, together with a detailed kinetic characterization of the motor properties, qualifies nonmuscle myosin-2C as the ideal human myosin to decipher the structure-function relationships in the myosin motor domain that underlie its kinetic signatures (4, 9).

To understand how structural adaptations lead to characteristic kinetic features, we solved the 2.25 Å crystal structure of the human nonmuscle myosin-2C motor domain (NM2C) (**Figure 1B**, **Table 1**). The NM2C pre-powerstroke state structure suggests that structural fine-tuning at the active site and a reduced interdomain connectivity in the motor domain define its kinetic signatures. Comparative structural analysis, ensemble solution kinetic studies, and *in silico* molecular dynamics (MD) simulations collectively demonstrate a communication pathway that connects the active site and the distal end of the motor domain. Disruption of the pathway uncouples the myosin ATPase activity from actin-activation, alters the ATP/ADP sensitivity and results in the loss of nonmuscle myosin-2 kinetic signatures. The allosteric coupling pathway may be of significance in intermolecular gating and load-sensitivity of nonmuscle myosins-2 that is important for their physiological function as cytoskeletal strain-sensor in the context of stress fibers, actin arcs, and cortical actin-based structures.

**Table 1.** Crystallographic data collection and refinement statistics for human NM2C.

| Parameter | Hs NM2C-ADP·VO<sub>4</sub> |
| --- | --- |
| *X-ray data reduction statistics* |  |
| Space group | *P*22<sub>1</sub>2<sub>1</sub> |
| *Unit cell dimensions* |  |
| *a*, *b*, *c* | 81.12 Å, 125.61 Å, 153.96 Å |
| $\alpha$, $\beta$, $\gamma$ | 90°, 90°, 90° |
| Resolution | 97.33 Å – 2.25 Å |
| Last shell | 2.35 Å – 2.25 Å |
| Total measurements | 1116790 |
| No. of unique reflections | 75357 |
| Last shell | 9069 |
| Wavelength | 0.91841 Å |
| *R*<sub>merge</sub> | 0.12 |
| Last shell | 0.23 |
| *I*/$\sigma$(*I*) | 13.6 |
| Last shell | 2.63 |
| Completeness | 0.998 |
| Last shell | 1.00 |
| Multiplicity | 14.82 |
| Last shell | 14.97 |
| *Refinement statistics* |  |
| Resolution | 34.77 Å–2.25 Å |
| Last shell | 2.31 Å–2.25 Å |
| No. of reflections (working set) | 71459 |
| No. of reflections (test set) | 3786 |
| *R*-factor | 0.2210 |
| Last shell | 0.2910 |
| *R*-free | 0.2410 |
| Last shell | 0.2920 |
| *No. of non-hydrogen atoms* |  |
| Macromolecules | 7697 |
| Ligands | 33 |
| solvent | 666 |
| *Average B-factor* |  |
| Protein | 76.06 Å<sup>2</sup> |
| Ligands | 35.59 Å<sup>2</sup> |
| Solvent | 63.37 Å<sup>2</sup> |
| *RMS deviations from ideal values* |  |
| Bond lengths | 0.01 Å |
| Bond angles | 1.16° |
| Ramachandran favored | 96.22 % |
| Ramachandran allowed | 3.78 % |
| Ramachandran outliers | 0 |

## Results

### Overall topology of pre-powerstroke NM2C

We determined a 2.25 Å pre-powerstroke state structure of human NM2C, one of the slowest myosins-2 characterized. Crystallization was achieved after truncation of the flexible N-terminal 45 residues and the fusion of the NM2C C-terminus to *Dictyostelium* α-actinin tandem repeats 1 and 2 (**Figure 1B**, **Figure 1 - Figure supplement 1B**, **Table 1**). This approach has been used successfully in prior structural and kinetic studies on NM2C and allows the detailed analysis of structure-function relationships in the myosin motor domain (4, 9).

The overall topology of NM2C shares the structural four-domain architecture of myosin-2 motor domains comprising the N-terminal subdomain (Nter), the upper 50 kDa subdomain (U50 kDa), the lower 50 kDa subdomain (L50 kDa), and the converter that terminates in the lever (**Figure 1C**) (10). The nucleotide-binding active site is formed by structural elements of the U50 kDa and allosterically communicates with the actin binding interface that is formed by distant structural elements of the U50 kDa and the L50 kDa. The NM2C C_α_ atoms superimpose with a root mean square deviation *(r.m.s.d.)* of 0.57 Å, 0.78 Å, and 0.73 Å to the closely related motor domain structures from chicken smooth muscle myosin-2 (PDB entry 1BR2), scallop striated muscle myosin-2 (PDB entry 1QVI), and *Dictyostelium* nonmuscle myosin-2 (PDB entry 2XEL) (**Figure 1 - Figure supplement 1C**) in the pre-powerstroke state.

NM2C pre-powerstroke state characteristics include a closed conformation of the nucleotide binding motifs switch-1 and -2 in the active site (**Figure 1B**). In this conformation, the characteristic salt bridge between switch-1 R261 and switch-2 E483 that is pivotal for the hydrolysis of ATP is formed (10-12). The relay helix is in a kinked conformation, the central seven-stranded β-sheet is untwisted, and the converter domain and the adjacent lever arm in the up-position (**Figure 1B**). NM2C switch-1 and lever dihedral φ, ψ angles are substantially changed when compared to the pre-powerstroke state structures of chicken smooth muscle myosin-2 (PDB entry 1BR2), indicating that small arrangements in the active site are coupled to conformational changes at the distal end of the motor domain (**Figure 1 - Figure supplement 1C-D**, **Supplementary table 1**). Concomitantly, the relative orientation of converter and lever arm deviates by ~8-10° when compared to the high resolution pre-powerstroke state crystal structure of chicken smooth muscle myosin-2 (PDB entry 1BR2) and resembled the orientation of the respective regions of scallop striated muscle myosin-2 (PDB entries 1QVI, 1DFL) motor domain structures (**Figure 1 - Figure supplement 1D**).

### Unique structural rearrangements in the NM2C active site in the pre-powerstroke state

Switch-1, switch-2, P-loop, and the purine-binding A-loop are the prototypic nucleotide binding motifs in the myosin active site that undergo conformational changes in response to nucleotide binding and release. The active site is flanked by loop-3 and a loop that connects helices J and K in the U50 kDa, hereafter referred to as JK-loop (**Figure 2A**). The JK-loop constitutes a major connection between switch-1 and the nucleotide is of great functional significance as a hotspot for human cardiomyopathy-causing mutations in cardiac myosin-2 and the site of exon 7 in *Drosophila* muscle myosin-2 (**Figure 2 - Figure supplement 1A**) (13-15).

**Figure 2:**
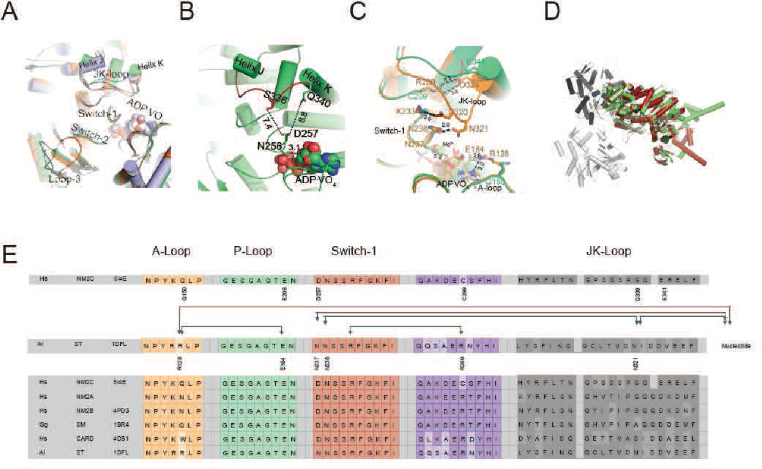
Conformational changes of the JK-loop in the myosin active site. (**A**) Top view on the NM2C active site in the pre-powerstroke state (green) superimposed on pre-powerstroke state structures form chicken smooth muscle myosin-2 (grey, PDB entry 1BR4), *Dictyostelium* nonmuscle myosin-2 (blue, PDB entry 2XEL), and scallop striated muscle myosin-2 (orange, PDB entry 1QVI). The nucleotide is shown in spheres representation. **(B)** Conformation of the JK-loop in vicinity to the NM2C active site. The JK-loop flanks the active site and connects helices J and K. The distance between the residue Q340 of the JK-loop in the U50 kDa and the D257 of switch-1 of the active site is ~8.8 Å. The distance between residue S336 of the JK-loop and switch-1 D257 is ~7.4 Å. Switch-1 residue N256 interacts with α-phosphate (3.1 Å) and β-phosphate (3.5 Å) group of ADP·VO_4_ in the active site. NM2C is colored in green/orange, the JK-loop is colored in brick red and ADP·VO_4_ is shown in spheres. **(C)** Interactions between the JK-loop and the switch-1 region are compared between the NM2C (green) and scallop striated muscle myosin-2 (orange, PDB entry 1QVI). A-loop residue R128 is coordinating the interaction to the ADP adenosine in the active site of striated muscle myosin-2. The distance between the residues is 3.2 Å. R128 further forms a hydrogen bond (2.8 Å) with E184 of the P-loop. JK-loop N321 is in hydrogen bond interaction with switch-1 N238, located at a distance of 4.6 Å to the hydroxyl group of the C2’ of the ADP ribose. The connectivity between switch-1 and the nucleotide is further strengthened by a hydrogen bond between N237 and the ADP ribose. NM2C lacks all interactions described for scallop striated muscle myosin-2 due to the replacement of R128 with Q150 and JK-loop shortening which increases the distance to the adenosine in the active site to 5.8 Å and disrupts constrains between swich-1 and the JK-loop. All residues in the JK-loop region are labeled for scallop striated muscle myosin-2 (PDB entry 1QVI). For NM2C only amino acid substitutions are labeled for legibility **(D)** Superimposition of the NM2C pre-powerstroke state structure (green) and the actin-bound near-rigor actoNM2C complex (red) shows that the nucleotide binding site does not undergo major structural changes. Actin subunits are colored in shades of grey and the nucleotide in the in spheres representation. **(E)** Sequence alignment of select structural elements in the myosin motor domain that interact with the JK-loop. Interactions are of A-loop R128 are highlighted with brackets for scallop striated muscle myosin-2 (PDB entry 1QVI). All highlights interactions are absent in NM2C due to the presence of Q150 in the A-loop. Abbreviations used: *Hs* NM2C: human NM2C (NP_079005.3); *Hs* NM2A: human nonmuscle myosin-2A (NP_002464.1); NM2B: human nonmuscle myosin-2B (NP_005955.3); *Gg* SM: chicken smooth muscle myosin-2 (NP_990605.2); *Hs* CARD: human beta β-cardiac muscle myosin-2 (NP_000248.2); *Ai* ST: scallop striated muscle myosin-2 (P24733.1). PDB entries are indicated when available.

A slight reduction in the JK-loop length in NM2C causes an 8.8 Å shift between switch-1 and the U50 kDa and abolishes the formation of a tight interaction network with residues of the active site found in other class-2 myosins (**Figure 2B,C**, **Figure 2 - Figure supplement 1A,B,C**). Comparative analysis of interactions between the JK-loop and the switch-1 region in NM2C and scallop striated muscle myosin-2 (PDB entry 1QVI) shows that in the latter, JK-loop N321 is in hydrogen bond interaction with switch-1 N238 and located at a distance of 4.6 Å to the hydroxyl group of the C2’ of the ADP ribose. The connectivity between switch-1 and the nucleotide is further strengthened by a hydrogen bond between N237 and the ADP ribose. Moreover, A-loop residue R128 forms hydrogen bonds with the ADP adenosine (3.2 Å) and E184 (2.8 Å) of the P-loop in the active site of striated muscle myosin-2. Strikingly, NM2C lacks all interactions described for scallop striated muscle myosin-2 due to the replacement of R128 with Q150 and the above mentioned JK-loop shortening. Both structural alterations increase the distance to the adenosine in the active site to 5.8 Å and disrupt constrains between swich-1 and the JK-loop (**Figure 2B,C,E**). The shift also increases the volume and hence the accessibility of the NM2C active site when compared to the transition state structures of chicken smooth muscle myosin-2 (PDB entry 1BR2), *Dictyostelium* nonmuscle myosin-2 (PDB entry 2XEL), and scallop striated muscle myosin-2 (PDB entry 1DFL,1QVI) (**Figure 2A**, **Figure 2 - Figure supplement 1B,D,E**). A 4.3-fold volume increase from 697 Å^3^ to 3054 Å^3^ in the active site is observed for example between scallop striated muscle myosin-2 (PDB entry 1QVI) and NM2C (**Figure 2 - Figure supplement 1D,E**). Strikingly, the enlarged accessibility of the active site region observed in the NM2C pre-powerstroke state structure resembles the conformation of the active site found in the actin-bound NM2C rigor state (PDB entry 5J1H), indicating that F-actin binding does not induce major structural rearrangements in the vicinity of the active site (**Figure 2C**).

The temperature factors obtained from the refined crystal structure show that the NM2C JK-loop is highly flexible despite its reduced length. We attribute the flexibility to the lack of constraints with switch-1 residues and the bound nucleotide (**Figure 2A,B,C,E**). Specifically, the connectivity of the JK-loop to switch-1 is reduced by the replacement of a conserved arginine (R280 in PDB entry 1QVI) with cysteine (C299) in the loop preceding helix I. This arginine further establishes an interaction with the highly conserved switch-1 K233 in scallop striated muscle myosin-2. The arginine to cysteine substitution in NM2C also abolishes the formation of a salt bridge with JK-loop residue E341 and hence disrupts the connection between the JK-loop and switch-1 (**Figure 2C,E**, **Figure 2 - Figure supplement 1C**). Consequently, the JK-loop loses its ability to sense conformational changes in the nucleotide binding motif switch-1 in response to ATP binding, hydrolysis, and product release.

The interaction between the JK-loop and the nucleotide is weakened by the replacement of an asparagine with G339, which abolishes hydrogen bond formation with switch-1 D257. Comparison with data from previous crystallographic studies shows that D257 replaces an asparagine (N238 in PDB entry 1QVI) in the switch-1 in striated muscle myosins-2. The asparagine interacts weakly with the ADP moiety in the pre-powerstroke state (**Figure 2B,D,E**, **Figure 2 figure supplement 1B,C**) and strongly in the near-rigor state (16, 17). The nucleotide coordination in NM2C is further weakened by residue Q150 that replaces an arginine (R128 in PDB entry 1QVI) in the A-loop of scallop striated muscle myosin-2. The substitution disrupts coordinating interactions between the A-loop and the ADP adenosine (**Figure 2D**, **Figure 2D**, **figure 2 supplement 1B,C**). The replacement additionally impairs the formation of a salt bridge with the invariant E206 (E184 in PDB entry 1QVI) at the distal end of the P-loop that is predicted to be involved in nucleotide recruitment to the active site (**Figure 2D, Supplementary table 2**) (16).

Previous studies established that the number of hydrogen bond interactions between P-loop, switch-1, and the Nter positively correlate with the thermodynamic coupling (**K_AD_/K**_D_), the efficiency of F-actin to displace ADP during the catalytic cycle (16). Substitution of an arginine with K255 and a glutamate with H700 in NM2C is expected to weaken the tight and highly conserved interaction network and is expected to contribute to the low thermodynamic (**K_AD_/K**_D_) and kinetic (**k_−AD_/k**_−D_) coupling efficiency, and the slow actin-activated ADP release in nonmuscle and smooth muscle myosins-2 compared to cardiac and striated muscle myosins-2 (**Supplementary table 2**) (16).

Taken together, comparative structural analysis of the active site of myosins-2 suggests that the interconnectivity of the highly conserved nucleotide switches and myosin subdomains is weaker in NM2C and other nonmuscle myosins-2 compared to fast sarcomeric myosins-2.

### Interdomain connectivity between converter, Nter, and lever

The interdomain connectivity between the converter and the Nter in the myosin-2 motor domain is established by the relay and the SH1-SH2 helix. The Nter controls the movement of the converter, which undergoes a large-scale rotation that drives the powerstroke during force generation (18-22).

Residue R788 at the interface of converter and lever is highly conserved in nonmuscle and smooth muscle myosins-2 and the only connecting hub between structural elements of the L50 kDa and the Nter (**Figure 3A,B**). Notably, replacement of R788 with a lysine in muscle myosins-2 weakens the interaction between the aforementioned structural elements in all states of the myosin and actomyosin kinetic cycle (**Figure 3 - Figure supplement 1A,B**). In NM2C, R788 forms 3 main chain and 5 side chain interactions with residues of the SH1-SH2 helix, the converter, and the lever in the pre-powerstroke state. Specifically, the guanidinium group of the R788 side chain of NM2C forms hydrogen bonds with the main chain carbonyls of residues Q730 and G731 of the SH1 helix. The hydrophobic methylene groups of R788 are stabilized by SH1 helix F732 and the main chain oxygen and hydroxyl groups of N776 of the converter. Y518 from the relay helix interacts with residues G731 and F732 of the converter, thereby interlinking R788 with both structural elements. The side chain of R788 is in van der Waals distance (4.8 Å) from W525 of the relay loop. The main chain carbonyl of R788 further interacts with V791 at the converter/lever junction (**Figure 3A,B**). This interaction is important for the stabilization of the converter fold and the interface with the lever in the absence of F-actin. Notably, the R788 side chain-main chain interaction is also evident in crystal structures of nonmuscle myosins-2 in the nucleotide-free (PDB entry 4PD3) and the phosphate-release state (PDB entry 4PJK) (not shown). We therefore hypothesize that (i) the tight interaction network formed by R788 is required for the precise positioning and coupling of the converter, the SH1-SH2 helix, the relay helix, and the lever arm throughout all steps of the myosin ATPase cycle (**Figure 1 - Figure supplement 1A**), and (ii) different communication pathways between the active site and the distal end of the motor domain are employed by NM2C in the presence and absence of F-actin.

**Figure 3:**
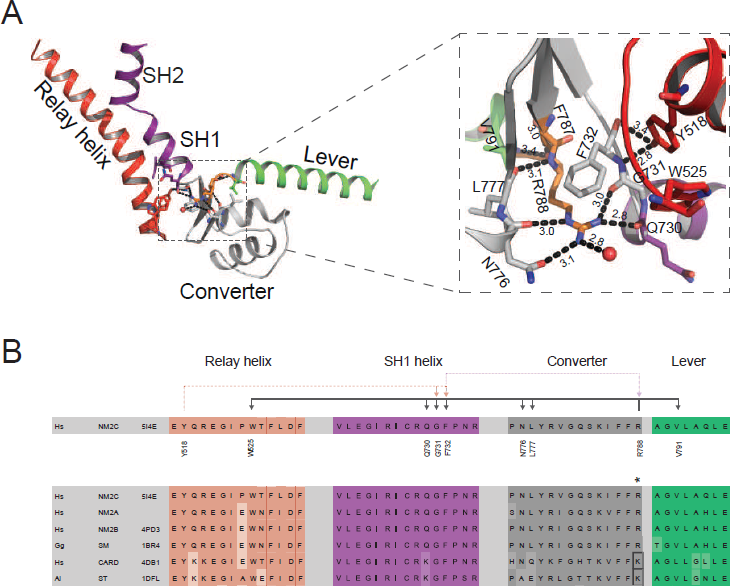
Interdomain connectivity at the converter/Nter/lever junction. (**A**) Interaction profile of R788 in the pre-powerstroke state. R788 is shown in orange colored sticks and the converter is colored in white, the relay helix in red, the SH1-SH2 helix in purple, and the lever arm in green colored cartoon representation. The inset shows a close-up view of the complete R788 interaction profile and is rotated 137° respective to the main panel. The guanidinium group of R788 forms hydrogen bonds (2.8 Å) with backbone oxygen atom of Q730 from the SH1 helix and the backbone oxygen atom of G731 (3.0 Å) of the converter. The δ-nitrogen atom of R788 interacts (3.0 Å) with N776 backbone oxygen atom of N776. The R788 guanidinium group interacts (3.1 Å) with the hydroxyl group of N776 of the converter. The backbone nitrogen atom of R788 interacts (3.1 Å) with the carbonyl group of L777 of the converter. The backbone carbonyl group of R788 interacts (3.4 Å) with the backbone nitrogen of V791 of the lever as well as a water molecule (3.0 Å). The hydroxyl group from relay helix Y518 interacts with the backbone nitrogen atom of G731 (2.8 Å) and backbone oxygen atom of F732 (3.4 Å). F732 forms hydrophobic interactions with the methylene groups of R788 with the latter positioned in van der Waals distance to relay loop W525. All the amino acids involved in interactions with R788 are shown as sticks and water molecules as spheres. **(B)** Sequence alignment of selected regions from relay helix, SH1 helix, converter and lever arm shows the high sequence conservation within the myosin-2 motor domain. The asterisk indicates the invariant, positively charged residue corresponding to NM2C R788. The interactions of NM2C R788 with structural elements of the L50 kDa, the converter, and the lever are highlighted. Abbreviations used: *Hs* NM2C: human NM2C (NP_079005.3); *Hs* NM2A: human nonmuscle myosin-2A (NP_002464.1); NM2B: human nonmuscle myosin-2B (NP_005955.3); *Gg* SM: chicken smooth muscle myosin-2 (NP_990605.2); *Hs* CARD: human beta β-cardiac muscle myosin-2 (NP_000248.2); *Ai* ST: scallop striated muscle myosin-2 (P24733.1). PDB entries are indicated when available. Lysine residues that replace R788 in cardiac and striated muscle myosins-2 highlighted in the boxed area.

### Kinetic consequences after the disruption of the converter/Nter/lever interface

To experimentally probe for a possible effect of R788 on myosin motor function, we performed comparative ensemble solution kinetic studies with NM2C and R788E, a mutant in which a glutamate replaces the converter R788. Rationale for the design of the R788E charge reversal mutant was to disrupt side chain:side chain and side chain:main chain interactions, an approach that has been successfully used in *in vitro* and *in vivo* studies to probe for the influence of the converter/U50 kDa interface on myosin-2 performance (19, 23-25).

R788E shows prominent changes in virtually all parameters of the myosin and actomyosin ATPase cycle (**Supplementary table 3**). Confirming our hypothesis, transient kinetic changes mainly affect nucleotide binding and release kinetics and are more pronounced in the absence of F-actin, as seen in single-turnover measurements (**Figure 4A,B**). Most importantly, nonmuscle myosin-2 specific transient kinetic signatures such as a high **k+_AD_/K_1_k+_2_** ratio (**k_+AD_/K_1_k_+2_** ~ 2-20) are absent in R788E (**k_+AD_/K_1_k_+2_** ~ 0.26) due to a 20-fold acceleration of the second-order rate constant for ATP binding (**K_1_k+_2_**) by F-actin (**Table 2, Figure 4 - Figure supplement 1A-D**) (4, 26, 27). This feature is also described for conventional myosins-2 from cardiac and striated muscle that bind ATP and ADP with similar rates (**Supplementary table 2**) (28). Actin-activation of the ADP release ***(k*_−AD_** = 0.68±0.01 s^-1^ for NM2C and ***k*_−AD_** = 0.19±0.01 s^-1^ for R788E) results in a kinetic coupling constant ***k*_−AD_**/k_−D_ of ~ 2.5 for R788E, whereas neutral or negative kinetic coupling is a feature of NM2C (**k_−AD_**/k_−D_ =0.7) and other human nonmuscle myosins-2 (**Table 2**) (4, 26, 27). The changes in ADP binding and release kinetics of R788E result in a 10-fold increase in the ADP dissociation equilibrium constant **K_AD_** (**K_AD_** ~ 0.29 μM for NM2C and ~ 2.68 μM for R788E). The thermodynamic coupling (**K_AD_**/K_D_=5) for R788E is 42-times higher than for NM2C (**Supplementary table 3**). R788E displays an extraordinary slow ADP binding rate constant (**k+_AD_**=0.03±0.001 μM^-1^s^-1^) (**Table 2, Figure 4 - Figure supplement 1D**) in the presence of F-actin whereas the kinetic constants for the interaction between actomyosin and ATP, ***K*_1_*K*_+2_** and 1/**K_1_** are marginally affected when compared to NM2C (**Supplementary table 3**). The F-actin affinity in the absence and presence of ADP (**K_A_**, **K_DA_**) of NM2C and R788E is similar (**Supplementary table 3**) and F-actin can activate the steady-state ATPase activity of R788E to approximately half the *k_cat_* of NM2C *(k_cat_* = 0.2±0.01 s^-1^ for R788E and *k_cat_* = 0.37±0.02 s^-1^ of NM2C) (**Supplementary table 3**) under steady-state conditions. The duty ratio at an F-actin concentration of 190 μM increases from ~ 0.3 for NM2C to ~ 1 for R788E due to the decreased *k_cat_* and the decreased actin-activated ADP release rate **k_−AD_**. This feature makes **k_−AD_** likely to rate-limit the kinetic cycle of R788E, whereas the actin-activated P_i_ release is expected to rate-limit the kinetic cycle of NM2C and other myosins-2 (1, 4, 26, 27, 29, 30).

**Figure 4:**
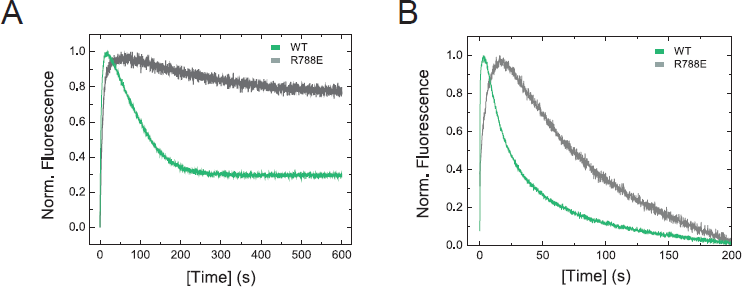
Transient kinetic features of NM2C and R788E. (**A,B**) Interaction between NM2C/R788E with ATP under single-turnover conditions in the absence **(A)** and presence **(B)** of F-actin. Binding 0.375 μM mantATP to 0.5 μM myosin/actomyosin results in a transient fluorescence increase that is followed by a short plateau (hydrolysis) and a slow decrease in mantADP fluorescence that is associated with its release. All three phases are reduced in R788E (grey) compared to NM2C (green) and are more pronounced in the absence of F-actin **(A)**. The very slow decrease of the fluorescence signal in R788E in **(A)** indicates that either the ATP hydrolysis rate or a subsequent release rate of the hydrolysis products are severely decreased when compared to NM2C.

**Table 2.** Comparative analysis of kinetic signatures of monomeric myosin-2 motor domain constructs. Abbreviations used: NM2A: human nonmuscle myosin-2A, NM2B: human nonmuscle myosin-2B (PDB entry 4PD3); NM2C: human nonmuscle myosin-2C; SM: chicken smooth muscle myosin-2 (PDB entry 1BR2); CARD: human/chicken β-cardiac myosin-2 (PDB entry 4DB1); ST: rabbit/scallop striated muscle myosin-2 (PDB entry 1DFL).

| Myosin | $k_{cat}/K_{app}$ | $k_{+AD}/K_1k_{+2}$ | $K_{AD}/K_D$ | $k_{-AD}/k_{-D}$ | $K_{DA}/K_A$ | duty ratio |
| --- | --- | --- | --- | --- | --- | --- |
| *Hs* NM2C | 0.003 | 2.1 | 0.12 | 1 | 0.13 | $\sim 0.3^{\$}$ |
| *Hs* R788E | 0.004 | 0.26 | 5 | 3 | 0.19 | $\sim 1^{\$}$ |
| *Hs* NM2A (26) | 0.002 | 19.4 | 0.7 | 2.8 | 2 | 0.1 |
| *Hs* NM2B (27) | 0.002 | 10 | 0.2 | 0.7 | 0.4 | 0.37 |
| *Gg* SM (28, 54) | 0.07 | 4.4 | 4.2 | 12 | 6.9 | <0.05 |
| *Hs/Gg* CARD (28, 55) | 0.07<sup>#</sup> | 1.6 | 42 | 103.3 | 23.9 | <0.02 |
| *Oc/Ai* ST (30, 56-58) | 1.6<sup>*</sup> | 1.6<sup>*</sup> | 49 | 250 | 30<sup>*</sup> | <0.04 |

\* Rabbit striated muscle myosin-2

The slow nucleotide binding and release kinetics of R788E suggest that the R788-mediated interaction at the converter/Nter/lever interface is allosterically communicated to the active site. This is in line with the observation that R788E does not show a nucleotide-induced change in the intrinsic tryptophan fluorescence signal during steady-state and transient-state kinetic assays (**Figure 5D,F, Figure 4 - Figure supplement 1E, Table 3**). The fluorescence change observed with NM2C is attributed to a conformation-induced change in the microenvironment of the conserved relay loop W525, which is in van der Waals distance to R788 and a direct indicator for the switch-2 induced converter rotation (31).

**Figure 5:**
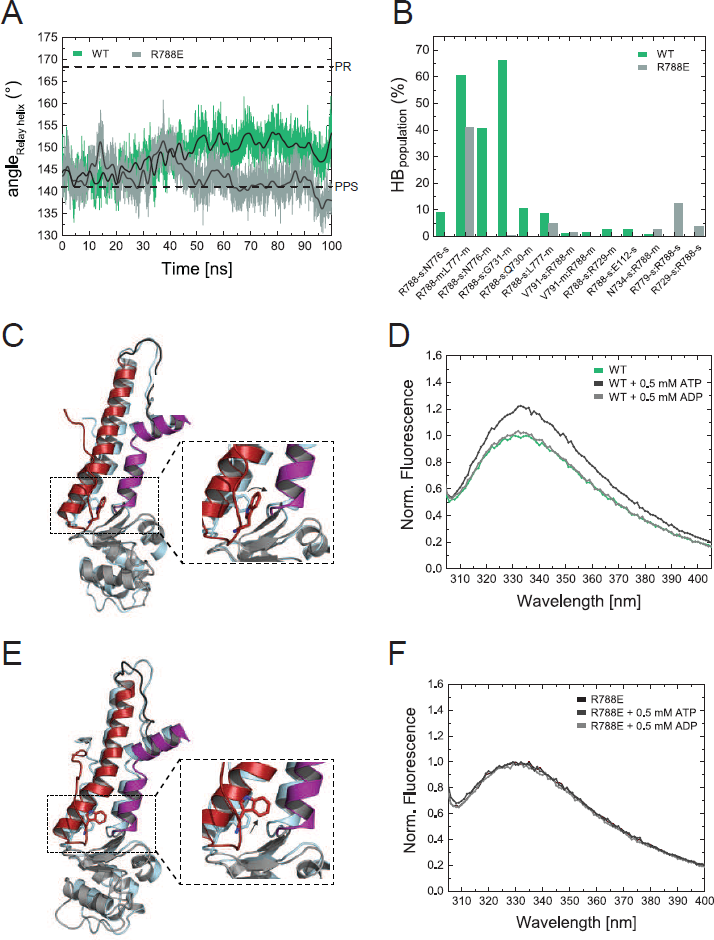
Structural importance of R788 at the converter/Nter/lever junction. (**A**) Relay helix angle as a function of MD simulation time, as monitored by the angle between C_α_ atoms of residues S489, M510, and E521 along the trajectories. The relay helix straightens in NM2C with a steady increase in the angle of the relay helix from approximately 145° to 150°, while the angle does not change significantly in R788E and fluctuates around 145° throughout the 100 ns time course of the simulation. Values for the relay helix angle observed in crystal structures of pre-power stroke (PPS) and post-rigor (PR) are indicated by dotted lines. **(B)** Population of hydrogen bonds (HB) between R788 (NM2C) or E788 (R788E) and surrounding structural elements over the simulation time of 100 ns. The abbreviations s and m indicate side chain and main chain. **(C, E)** Direct comparison of the dynamics and conformational changes in NM2C **(C)** and R788E **(E)** during MD simulations. Snapshots from the start (0 ns simulation time) and end conformations (100 ns simulation time) are shown in light cyan and colored cartoon representation, respectively. The relay helix is shown in red, the SH1-SH2 helix in purple and the converter in grey. Relay loop W525 is shown in red in stick representation. The insets show a close-up view of the conformational changes of W525 along the simulation trajectory. **(D, F)** Tryptophan fluorescence emission spectrum of 4 μM NM2C **(D)** and 4 μM R788E **(F)** in the absence of nucleotide or the presence of 0.5 mM ATP or 0.5 mM ADP.

### Allosteric communication pathway between the active site and the distal end of the motor domain

R788 is a hub amino acid and forms the center of a cluster of interactions that connect the converter, the SH1-SH2 helix, the relay helix, and the lever (**Figure 3A,B**). To understand its dynamic interactions that are allosterically communicated to the active site, we performed comparative molecular dynamics simulations of NM2C and R788E in explicit water with Mg^2+^. ATP bound to the active site over a time of 100 ns. As expected, the characteristic salt bridge between switch-1 R261 and switch-2 E483 of the active site that is critical for the hydrolysis of ATP is stable throughout the time course of the simulation for NM2C (**Figure 5 - Figure supplement 1A**). The relay helix that connects the active site and the converter straightens and likewise the converter undergoes a 27° rotation that directs lever arm motion (**Figure 5A,C**). This straightening is caused by the exchange of the co-crystallized ADP-VO_4_ to ATP, since the replacement with ADP did not lead to either relay straightening nor converter rotation in a control simulation. Hence, ATP appears to trigger the structural transition from the pre-powerstroke state (up position) towards the post-rigor state (down position). After 100 ns, the position of the converter and the adjacent lever show an orientation between the post-rigor and pre-powerstroke state, as compared to the crystal structures of *Dictyostelium* nonmuscle myosin-2 motor domain (PDB entries 1FMW, 2XEL) (**Figure 5C**). With the movement of the relay helix, the indole ring of relay loop W525 changes its conformation by 70-80°, which agrees with the experimentally observed nucleotide-induced change in the intrinsic tryptophan fluorescence signal in NM2C (**Figure 5B,C, Figure 4 - Figure supplement 1E**). The hub amino acid R788 is in transient interactions with main chain and side chain atoms of 10 amino acids of the SH1 helix and the converter during the 100 ns time course of the simulation (**Figure 5B**).

MD simulations for R788E indicate that all interdomain interactions between R788E of the converter and the SH1 helix are lost (**Figure 5B**). The straightening angle of the relay helix remains constant and abolishes a detectable converter rotation (**Figure 5A,E**). This allosteric decoupling at the distal end of the myosin motor domain is translated further upstream via the relay helix and results in a pronounced 6 Å conformational change of a loop that connects the γ-phosphate sensor switch-2 of the active site and the relay helix. As a consequence, the salt bridge between switch-2 E483 and switch-1 R261 appears less stable in the MD simulations of R788E as compared to NM2C (**Figure 5 - Figure supplement 1A,B**). The importance of the salt bridge between both switches for the ATP hydrolysis is established and expected to directly contribute to the experimentally observed impaired nucleotide binding, hydrolysis, and release kinetics of R788E in transient-state kinetic assays (**Supplementary table 3, Figure 4A, Figure 4 - Figure supplement 1A,D**) (11, 32, 33). Moreover, the side chain of switch-1 N256 changes its orientation, thereby impairing hydrogen bond interactions with the α- and β-phosphate moieties of the nucleotide and constrains of switch-1. The lack of coordinating interactions is reflected in the very slow nucleotide binding and release rate constants (**Figure 4A, Figure 4 - Figure supplement 1A, Supplementary table 3**).

The side chain of relay loop W525 does not undergo a conformational change throughout the time course of the simulation for R788E, supporting our experimental observation that R788E does not exhibit a nucleotide-induced change in its intrinsic tryptophan fluorescence signal of R788E (**Figure 5D,F, Figure 4 - Figure supplement 1E**). The direct comparison of the dynamic interaction and conformational signatures of W525 in NM2C and R788E reveals that R788 is in van der Waals distance (5.4 Å) from relay loop W525 at the start of the simulations. W525 transiently interacts with I523 (18%) and Q519 (14%), changes its conformation by 70-80° and increases the distance to NM2C R788 to 7.3 Å after 100 ns simulation time. In comparison, the distance between W525 and E788 increases to 9.9 Å during the simulation for R788E. The lack of a conformational change in W525 along the simulation trajectory results in transient interactions with Q515 (55%) and Q730 (14%) instead of the interactions with I532 and Q519 as seen in NM2C.

Taken together, our *in silico* and experimental *in vitro* data support a model in which R788 is a distant allosteric modulator of switch-2 dynamics at the active site that impacts nucleotide binding and release kinetics in the actin-detached states. Moreover, our data support a model of R788 as a quencher of W525 fluorescence.

## Discussion

The experiments presented here collectively demonstrate an allosteric communication pathway from the distal end of the myosin motor domain that, together with substitutions of several key residues in or in vicinity to the active site, account for NM2C-specific kinetic properties. Disruption of the pathway by mutation of R788 to a glutamate causes the loss of its enzymatic signatures and results in a high duty ratio motor.

It is of note that several key residues involved in the communication pathway are near residues that are mutated in patients with autosomal dominant hearing loss (34). Missense mutation G376C is in proximity to residues C324 of helix J and R328 of the JK-loop (**Figure 4 - Figure supplement 1F**). Based on its location, we suggest that this substitution may interfere with the nucleotide binding and release kinetics from the NM2C active site. Missense mutation R726S is in the SH1-SH2 helix and the guanidinium group of the wild type arginine interacts (3.3 Å) with the NM2C Nter. The serine residue is expected to disrupt this interaction because of the shorter side chain and different charge when compared to arginine. It is therefore likely that these mutations impact the interaction of the motor domain with nucleotides, thereby contributing to impaired tension-sensing and maintenance of nonmuscle myosin-2C in the human cochlea (34-36).

### R788 is part of a conserved pathway that connects the active site and the converter

R788 is part of an allosteric communication pathway that connects the converter at the distal end of the myosin motor domain via the relay helix with switch-2 of the active site. Uncoupling of the converter from the motor domain in R788E slows down all kinetic steps of the myosin ATPase cycle and decreases its actin-activation, due to altered switch-2 dynamics. The observed kinetic phenotype of the R788E myosin ATPase cycle is similar to the *Dictyostelium* nonmuscle myosin-2 non-hydrolyzer mutants in which the salt bridge between switch-1 and switch-2 is destroyed by mutagenesis (11, 32). Non-hydrolyzer mutants are characterized by long-lived ATP states and reduced nucleotide release and binding kinetics, including a drastic decrease in **k+_AD_** (11, 32). Like R788E, *Dictyostelium* myosin-2 non-hydrolyzer mutants do not exhibit a nucleotide-induced change in the intrinsic fluorescence signal (11). It is of note that the lack of an intrinsic fluorescence signal in R788E is caused by the blockage of the communication pathway on the relay helix before or at Y518 (**Figure 6A,B**) whereas the impairment of the nucleotide switches to form a salt bridge and the resulting lack of the switch-2 induced conformational change of the relay helix is the expected cause in the non-hydrolyzer mutants.

**Figure 6:**
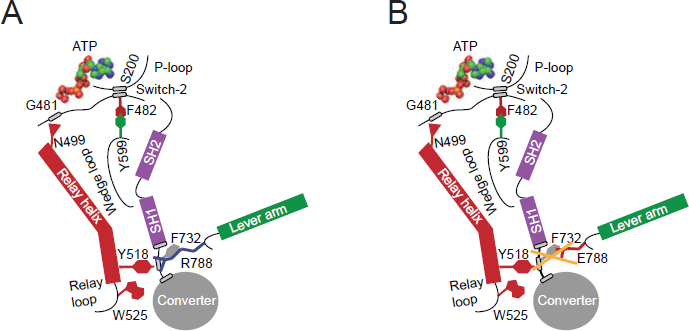
Proposed allosteric communication pathway from the converter to the NM2C active site. (**A**) Residue R788 connects the converter to the SH1 interface through main chain and side chain interactions in NM2C. This interface further interacts with Y518 of the relay helix and W525. The interactions are propagated to the active site and *vice versa* through the relay helix and through an interaction of relay helix residue N499 with the main chain of G481 from switch-2. The latter directly interacts with the nucleotide. Y599 of the wedge loop, that itself contacts the relay helix, establishes an interaction with switch-2 F482. This residue is in contact with S200 of the P-loop by main chain interactions and directly interacts with the nucleotide in the active site. **(B)** The interface between R788E and both, the SH1 and the relay helix, is disrupted. The allosteric communication from the active site to the converter is compromised. The experimental observation that R788E does not change its intrinsic fluorescence upon nucleotide binding which is caused by a lacking conformational change of W525 indicates that the communication pathway is interrupted in the relay helix. The position of W525 in the relay loop at the distal end of the relay helix indicates that the pathway is interrupted before or at Y518, which is supported by the observation that the relay loop does not change its position during the time course of the MD simulation. As a consequence, Y518 cannot establish an interface with E788 and F732 of the converter and SH1 helix. Further, E788 cannot establish interactions with N776 and V791 and completely disrupts the structural integrity of the interface of converter, SH1-SH2 helix, relay helix and the lever arm and uncouples nucleotide-induced changes in the active site from the converter rotation. Only key residues involved are showed in the proposed mechanism. Small elliptical shapes show main chain interactions.

F-Actin affinities are largely unaffected by the R788E mutation, which is in line with the observation that switch-1 dynamics are only marginally affected in comparative MD simulations. The presence of F-actin however establishes ATP/ADP selectivity in the actomyosin ATPase cycle: ActoNM2C preferentially binds ADP over ATP, whereas actoR788E preferentially binds ATP over ADP (**Supplementary table 3**). The ATP selectivity is caused by a 85-fold decreased second-order ADP binding rate constant **k+_AD_**, a key signature of R788E actomyosin ATPase cycle. ATP/ADP selectivity is established by actin-induced conformational changes in the myosin motor domain and underlines that the coupling mechanism from the active site to the converter and *vice versa* is different in the presence and absence of F-actin. This observation is in line with recent reports on distinct pathways for the myosin and actin-activated ATPase cycle (21).

### Implications for NM2C function *in vivo*

Nonmuscle myosin-2C assembles into small bipolar filaments that dynamically tether actin filaments in cells (8, 37). The actomyosin interaction and hence the tension exerted by a sarcomeric array of nonmuscle myosin-2C is of importance for the function and organization of the apical junctional complex in the organ of Corti and actin-rich structures including stress fibers and actin arcs (8).

Based on our kinetic data, the calculated duty ratio of NM2C suggests that ~ 14 motor domains would be needed to be geometrically capable of interacting with F-actin at any given time. This number is identical to the number of motor domains per nonmuscle myosin-2C half filament, suggesting that the filament may be at the threshold of being processive in the absence of external loads (37). It is likely that this threshold is crossed in the presence of other actin binding proteins, intermolecular loads, and gating between the F-actin bound motor domains (38).

The structural prerequisites underlying gating and load-sensitivity in nonmuscle myosins-2 have not been investigated but include a distortion at the converter/lever or the converter/motor domain interface in the nanometer range that is allosterically communicated through the lever arm via the converter to the active site (39). Resisting load applied to the lever slows down the actin-activated ADP release from the lead motor of nonmuscle myosin-2A around 5-fold thereby increasing the duty ratio, but does not alter the rate of ATP binding (38, 39). The kinetic signatures of the strained nonmuscle myosin-2A lead motor are very similar to the observed kinetic features of R788E. The reduction of the steady-state ATPase of R788E and the concomitant increase in duty ratio and a rate-limiting ADP release rate goes in line with this finding. We propose that internal strain in the nonmuscle myosin-2 dimer distorts the lead motor at the converter/lever interface and leads to its axial translation as seen in electron microscopic studies (40). This translation abolishes the interaction of the converter R788 with residues of the SH1-SH2 helix and the relay helix, thereby uncoupling the myosin subdomains and disrupting the communication pathway from the converter to the active site, which is expected to result in a similar kinetic effect as in R788E. Consequently, a motor in a nonmuscle myosin-2 filament would decrease its enzymatic activity and stay strongly bound to F-actin. This feature is required to generate processivity and cytoskeletal tension and of physiological significance in the maintenance of cell shape and tensional homeostasis in the actin cytoskeleton.

## Material and Methods

### Protein production

For structural studies, a His_8_-tagged human NM2C construct comprising amino acids 45-799 directly fused to spectrin repeats 1 and 2 from α-actinin was generated based on the vector pFastBac1-NMHC-2C0-2R-His_8_ (4). Mutagenesis was accomplished by sequence-specific deletion using a whole-plasmid amplification approach. For kinetic studies, an equivalent motor domain construct comprising amino acids 1-799 of NM2C was cloned into a modified pFastBac1 vector containing a cDNA sequence encoding a C-terminal Flag-tag (**Figure 1 - Figure supplement 1B**). This construct was used as a template to introduce the R788E mutation by InFusion cloning (Clontech, Mountain View, CA 94943, USA). All proteins were recombinantly overproduced in the S*f*9/baculovirus system, purified to electrophoretic homogeneity, and concentrated to ~10 mg/ml using Vivaspin ultrafiltration units (Sartorius, Göttingen, Germany) as previously described (4, 41). Throughout this manuscript, numbering refers to the amino acid sequence of full-length nonmuscle myosin-2C (GenBank accession number NP_079005).

### Crystallization of NM2C

NM2C at a concentration of ~10 mg/ml was complexed with the ATP analogue ADP·VO_4_ and crystallized using the hanging drop vapor diffusion method by mixing 2 μl of protein solution with 2 μl of reservoir solution containing 50 mM Tris pH 8.2, 10 % (w/v) PEG-5K MME, 1 % (v/v) MPD, and 0.2 M NaCl at 8°C. Rectangular plate shaped crystals grew up to 400 × 300 × 400 μm^3^ within 4 weeks. Crystals were soaked in the corresponding mother liquor supplemented with 100 mM NaCl and 20 % (w/v) ethylene glycol. Protein crystals were transferred in serial steps of increasing concentrations of the cryo-solution. Crystals were transferred into liquid nitrogen using MiTeGen loops and stored at 100 K until data collection.

### Data collection, processing and refinement

X-ray diffraction data of NM2C crystals were collected at the beam line BL14.1 at Bessy II (Helmholtz-Zentrum, Berlin, Germany) to 2.25 Å resolution (**Table 1**). Data processing was performed using XDS and SADABS (42). The structure of NM2C in the pre-powerstroke state was solved by molecular replacement using Phaser (43). The crystal structure of the chicken smooth muscle myosin-2 motor domain (PDB entry 1BR4) was used as a search model and the two α-actinin repeats were manually traced and rebuilt. The electron density map was sharpened using Coot (44) to ensure the directionality and identity of the α-helices for the two α-actinin repeats. Maximum likelihood crystallographic refinement was performed using iterative refinement cycles in autoBUSTER (45). Iterative cycles of model building were performed using Coot, and model bias was minimized by building into composite omit maps. The model was initially validated in Coot and final validation was performed using MolProbity (46).

### Kinetic experiments

The actin-activated ATPase assays under steady-state conditions was performed as described earlier in buffer containing 10 mM MOPS pH 7.0, 50 mM NaCl, 2 mM MgCl_2_, 2 mM ATP, 0.15 mM EGTA, 40 U/ml l-lactic dehydrogenase, 200 U/ml pyruvate kinase, 200 μM NADH, and 1 mM phosphoenolpyruvate at 25°C with a Cary 60 Bio spectrophotometer (Agilent Technologies, Wilmington, DE 19808, USA) (41). Transient state kinetic assays were carried out as described previously with a TgK Hi-tech Scientific SF-61 DX stopped-flow system (TgK Hi-tech Scientific Ltd., Bradford-on-Avon, UK) in SF-buffer (25 mM MOPS pH 7.0, 100 mM KCl, 5 mM MgCfe and 0.1 mM EGTA) unless stated otherwise (4). Initial data fitting was performed with Kinetic Studio Version 2.28 (TgK Hi-tech Scientific Ltd., Bradford-on-Avon, UK). Plots were generated with OriginPro 8.5 (OriginLab Corp., Northampton, MA 01060, USA). Data interpretation is according to the kinetic scheme of the myosin and actomyosin ATPase cycle as presented in **Figure 1 - Figure supplement 1A**.

### Fluorescence measurements

Tryptophan fluorescence emission spectra of 4 μM NM2C or R788E in the presence and absence of 0.5 mM ADP and 0.5 mM ATP, respectively were measured after excitation at 297 nm at a temperature of 20 °C in a QuantaMaster fluorescence spectrophotometer (Photon Technology International, Birmingham, NJ 08011, USA) as described previously (47). Prior to the assay, proteins were transferred to SF-buffer with zebra spin desalting columns (Thermo Fisher Scientific GmbH, Dreieich, Germany). The fluorescence was normalized to the maximum fluorescence of NM2C or R788E in the absence of nucleotide.

### Molecular dynamics simulations

Molecular dynamics based *in silico* site directed mutagenesis and simulations were performed using NAMD 2.9 and the CHARMM27 force field (48, 49). The X-ray crystal structure of the NM2C motor domain in the pre-powerstroke state, encompassing residues 49-807, served as the starting structure for NM2C and R788E simulations. Mutations were introduced *in silico* and the proteins were prepared and optimized prior to MD simulations using the Protein Preparation Wizard of the Schrödinger software suite (Schrödinger Suite 2012 Protein Preparation Wizard; Epik version 2.3; Impact version 5.8; Prime version 3.1; Maestro version 9.3. Schrödinger LLC, New York, NY, USA). The proteins were fully hydrated with explicit solvent using the TIP3P water model and charge neutralization was accomplished by adding Na+ counter ions (50). Short-range cutoffs of 12 Å were used for the treatment of non-bonded interactions; while longrange electrostatics was treated with the particle-mesh Ewald method (51). All simulations were carried out in an NpT ensemble at 310 K and 1 atm using Langevin dynamics and the Langevin piston method. A 1 fs time step was applied. Prior to production runs the solvated systems were subjected to an initial energy minimization and subsequent equilibration of the entire system for 5 to 10 ns. All MD simulations were carried out at the Computer Cluster of the Norddeutscher Verbund für Hoch- und Höchstleistungsrechnen.

## Acknowledgements

We thank the staff at the beamline BL14-1 at BESSY for technical support. We thank the Norddeutscher Verbund für Hoch- und Höchstleistungsrechnen (HLRN) for providing computational resources and the Biophysics Core of the National Heart, Lung, and Blood Institute (NHLBI) for advice, support and the use of the facility. Data deposition: The atomic coordinates and structure factors have been deposited in the Protein Data Bank, www.pdb.org (PDB entry 5I4E).

## Author Contributions

D.J.M. conceived the study. K.C. performed the crystallization, collected the diffraction data, and solved the structure. S.M.H. cloned, expressed and purified the proteins, and performed the biochemical and kinetic assays. M.P. performed and analyzed the molecular dynamics simulations. K.C., S.M.H., M.P., J.R.S., and D.J.M. designed experimental approaches and wrote the manuscript. All authors read and approved the final manuscript.

## Competing Interests

The authors declare no conflicts of interest with the contents of this research article.

## Supplementary Information

**Figure 1 - Figure supplement 1:**
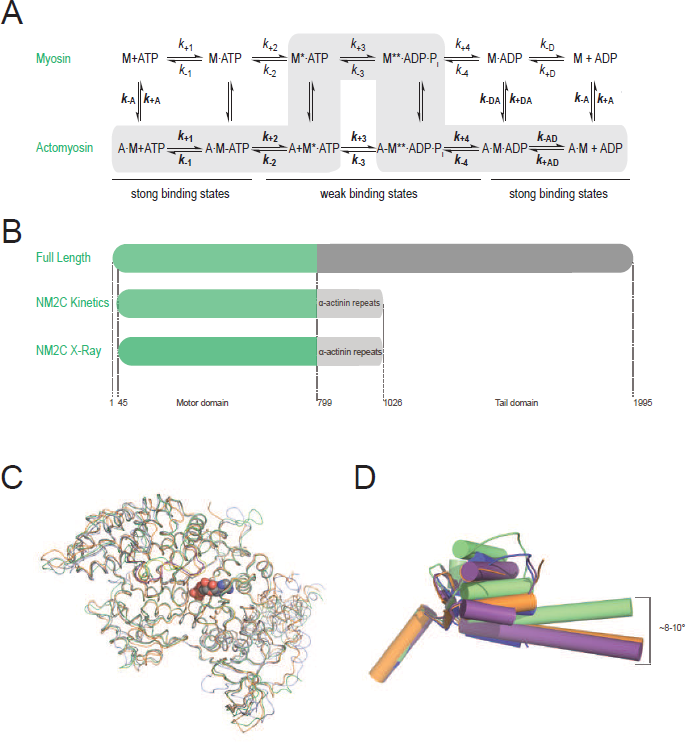
Myosin-2 ATPase cycle, expression constructs, and structural alignment of the NM2C C_α_ coordinates. (**A**) Consensus scheme of the myosin and actomyosin ATPase cycle. The upper part represents the myosin (M) ATPase cycle. The lower part represents the ATPase cycle in the presence of F-actin (A). The asterisk denotes enhanced states of the intrinsic myosin fluorescence, which are attributed to the nucleotide-induced changes in the microenvironment of the conserved relay loop W525. Lowercase *k* denotes a rate constant. *k*_+D_=*k*_−6_, *k*_−D_=*k*_−6_, ***k*+_AD_=k_−6_**, ***k*_−AD_=k_−6_**. An uppercase *K* denotes a dissociation equilibrium constant (*K*=*k*_*-x*_/*k*_+x_) throughout this work. Normal and bold face notation denote the respective kinetic constants in the absence and presence of F-actin. The main pathway of the actomyosin ATPase cycle is highlighted in grey. Strong and weak actin binding states are indicated. **(B)** Schematic representation of the expression constructs used in this study. Top, Nonmuscle myosin-2C contains a N-terminal motor domain (green) followed by a neck and a tail domain (dark grey). *Middle*, For kinetic studies of NM2C and R788E, the motor domain was directly fused to spectrin repeats 1 and 2 from *Dictyostelium* α-actinin (light grey) which serves as an artificial lever arm. The concept of the artificial lever arm has been successfully used in structural and kinetic studies of myosins-2 (4, 11, 59, 60). *Bottom*, For structural studies, the N-terminal 45 amino acids were deleted. The numbering refers to the amino acid sequence of the full-length protein. **(C)** The NM2C C_α_ atoms (green) in ribbon representation superimpose with a root mean square deviation *(r.m.s.d.)* of 0.57 Å to chicken smooth muscle myosin-2 (grey, PDB entry 1BR2), with 0.78 Å to scallop striated muscle myosin-2 (orange, PDB entry 1QVI), and 0.73 Å to *Dictyostelium* nonmuscle myosin-2 (violet, PDB entry 2XEL), underlining a strong correlation between C_α_ geometry and overall motor domain fold. The JK-loop and the nucleotide are highlighted in color and spheres representation. **(D)** Relative orientation of converter and lever in NM2C (green), chicken smooth muscle myosin-2 (PDB entry 1BR2, blue), and scallop striated muscle myosin-2 (PDB entry 1QVI (orange), 1DFL (violet)) motor domain structures in cartoon representation. α-helices are depicted as cylinders.

**Figure 2 - Figure supplement 1:**
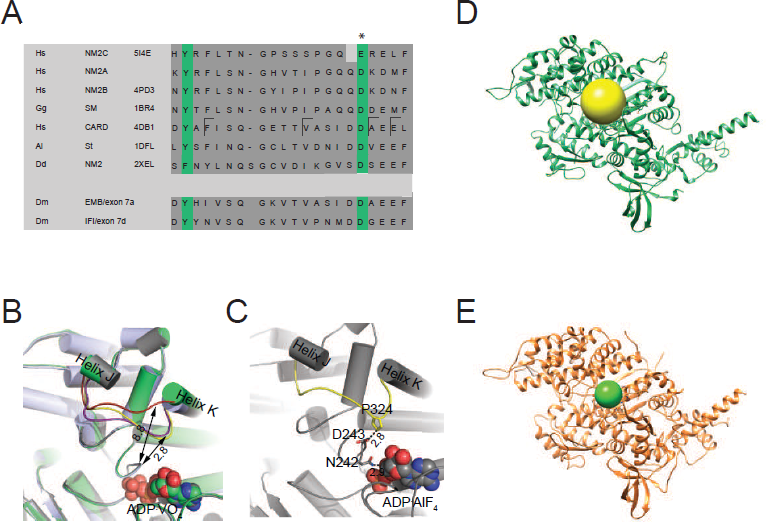
Active site characteristics in myosin-2 motor domains. (**A**) Sequence alignment of myosin-2 JK-loops. The asterisk indicates the invariant, negatively charged residues corresponding to NM2C E341. The abbreviations used are as follows: *Hs* NM2C: human NM2C (NP_079005.3); *Hs* NM2A: human nonmuscle myosin-2A (NP_002464.1); NM2B: human nonmuscle myosin-2B (NP_005955.3); *Gg* SM: chicken smooth muscle myosin-2 (NP_990605.2); *Hs* CARD: human beta β-cardiac muscle myosin-2 (NP_000248.2); *Ai* ST: scallop striated muscle myosin-2 (P24733.1); *Dd* NM2: *Dictyostelium* nonmuscle myosin-2: (XP_637740.1); *Dm* EMB: *Drosophila* embryonic body wall muscle myosin-2 (P05661 with spliced exon 7a); *Dm* IFI: *Drosophila* indirect flight muscle myosin-2 (P05661 with spliced exon 7d). PDB entries are indicated when available. Cardiomyopathy-associated mutations in β-cardiac myosin-2 on positions F312 (F312C), V320 (V320M), A326 (A326P), and E328 (E328G) are highlighted in the boxed areas. **(B)** Superimposition of JK-loops of pre-powerstroke state structures of NM2C (brick red), chicken smooth muscle myosin-2 (yellow, PDB entry 1BR2), and *Dictyostelium* nonmuscle myosin-2 (purple, PDB entry 2XEL) motor domains are shown in close-up view. The distance between the JK-loops of smooth muscle myosin-2 and *Dictyostelium* nonmuscle myosin-2 and switch-1 is ~ 2.8 Å and 8.8 Å for NM2C. **(C)** The distance between the residue P324 of the JK-loop (yellow) in the U50 kDa of chicken smooth muscle myosin 2 (grey, PDB entry 1BR2) and the D243 of switch-1 is ~2.8 Å. Switch-1 residue N242 interacts with α-phosphate (2.8 Å) and β-phosphate (3.1 Å) group of the ADP-ALF_4_ complex in the active site is shown in spheres. **(D,E)** The volume of the active site, indicated by the inclusion spheres, was determined to 3065 Å^3^ in NM2C **(D)** and 697 Å^3^ in scallop striated muscle myosin-2 (PDB entry 1QVI) **(E)**. Both structures are in the pre-powerstroke state.

**Figure 3 - Figure supplement 1:**
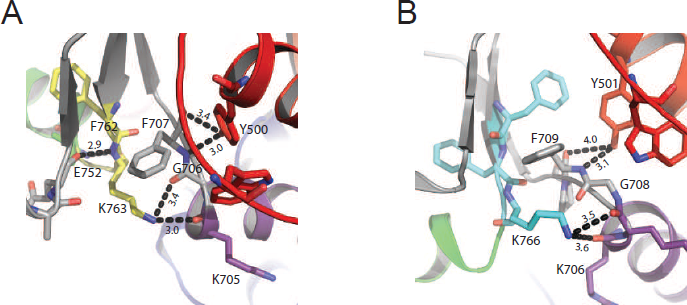
Interdomain connectivity at the converter/Nter/lever junction in muscle myosins-2. In contrast to NM2C (**Figure 3**), the substitution of R788 with a lysine K762 in scallop striated muscle myosin-2 (PDB entry IQVI) **(A)** and K766 in human β-cardiac muscle myosin-2 (PDB entry 4DB1) **(B)** reduces the number of side chain and main chain interactions at the converter/Nter/lever junction. Coloring is according to **Figure 3**.

**Figure 4 - Figure supplement 1:**
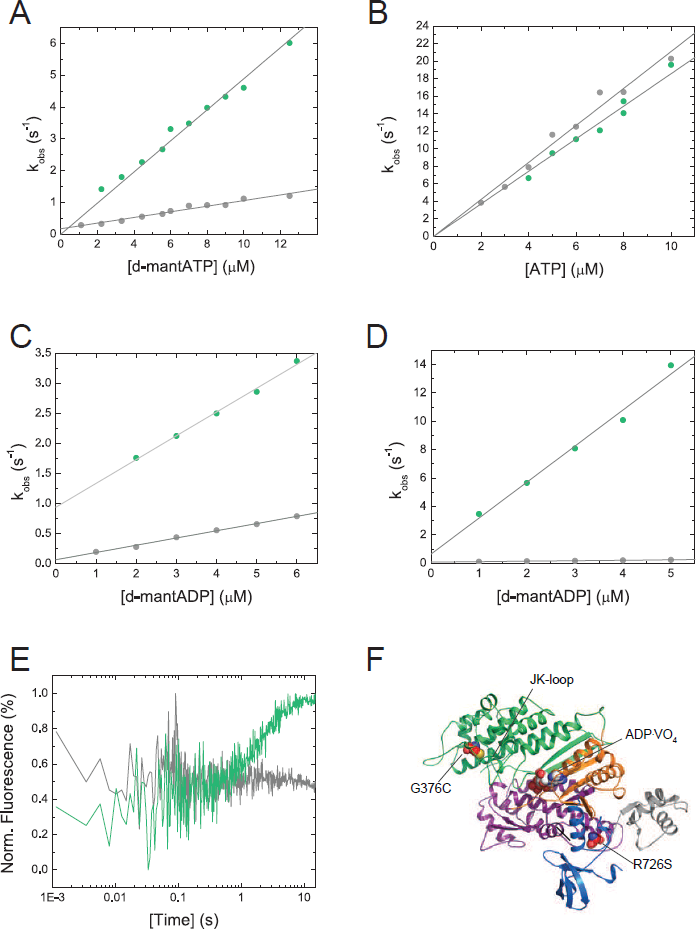
Nucleotide binding characteristics of NM2C and R788E and disease causing NM2C mutations. (**A**) Dependence of the observed rate constants (*k*_obs_) upon d-mantATP binding to 0.25 μM NM2C/R788E on the nucleotide concentration. Linear fits to the data result in second-order rate binding constants of *K*_1_*k*_+2_ = 0.48±0.01 μM^-1^s^-1^ for NM2C (green) and a 5-fold reduced binding rate constant of *K*_1_*k*_+2_ = 0.09±0.005 μM^-1^s^-1^ for R788E (grey). **(B)** The second-order ATP binding rate constant ***K*_1_*k*_+2_**, determined by the ATP-induced dissociation of the pyrene-labeled actoNM2C (green) or actoR788E (grey) complexes are with ***K*_1_*k*_+2_** =1.23±0.02 μM^-1^s^-1^ and ***K*_1_*k*+_2_** = 2.06±0.07 μM^-1^s^-1^ similar. **(C)** The R788E (grey) mutation decreases the second-order ADP binding rate constant 3-fold to *k*_+D_ = 0.12±0.01 μM^-1^s^-1^ when compared to k_+D_ = 0.39±0.01 μM^-1^s^-1^ for NM2C (green). **(D)** The ADP binding rate constant ***k*_+AD_** = 0.03±0.001 μM^-1^s^-1^, as calculated from the slope of the k_obs_ versus [d-mantADP] plot, is 85-fold decreased for R788E (grey) when compared to ***k*_+AD_** = 2.54±0.18 μM^-1^s^-1^ for NM2C (green). **(E)** Time-dependent change in the intrinsic tryptophan signal upon mixing 0.5 mM ATP with 0.25 μM NM2C (grey) or R778E (grey) in the presence of 5 μM ADP. ATP binding increases the fluorescence signal in NM2C but not R788E under identical conditions in a stopped-flow spectrophotometer. **(F)** Missense mutations G376C and R726S are associated with autosomal dominant hearing impairment (DFNA4) and their location in NM2C is shown in spheres representation. G376C is in proximity to the JK-loop, R726S is in the SH1-SH2 helix. NM2C subdomains are color coded according to **Figure 1C** and the nucleotide is shown in spheres representation.

**Figure 5 - Figure supplement 1:**
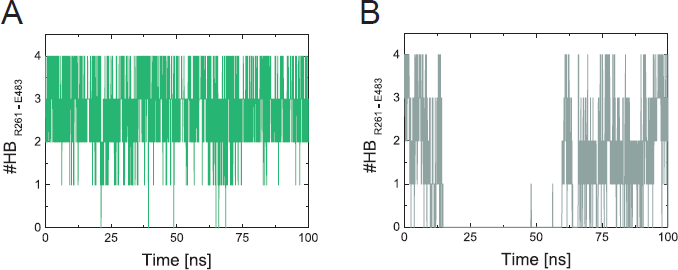
Stability of the salt bridge between switch-1 and switch-2 in MD simulations of NM2C and R788E. (**A,B**) Monitored number of hydrogen bond (#HB) interactions between R261 (switch-1) and E483 (switch-2) along the simulation time of NM2C **(A)** and R788E **(B)**. Hydrogen bonds were detected with cutoff values for the donor-acceptor distance and angle of 3.5 Å and 30°. Note the intermediate breaking of the salt-bridge in R788E **(B)**.

**Supplementary table 1:** Dihedral angles of switch-1 and lever arm residues in crystal structures of Nm2C and smooth muscle myosin-2 (PDB entry 1BR2).

| ***Hs* NM2C** |  |  | ***Gg* smooth muscle myosin-2** |  |  |
| --- | --- | --- | --- | --- | --- |
| Residue | $\varphi$ | $\psi$ | Residue | $\varphi$ | $\psi$ |
| *Switch-1* |  |  |  |  |  |
| N256 | -152.2 | 125.4 | N242 | -148.5 | 139.7 |
| D257 | -66.7 | -15.6 | D243 | -78.1 | -32.9 |
| N258 | -135.6 | 51.3 | N244 | -116.9 | 55.9 |
| S259 | -71.5 | 128.2 | S245 | -81.2 | 127.9 |
| S260 | -78.1 | 122.8 | S246 | -71.1 | 119.6 |
| R261 | -104.7 | 29.2 | R247 | -102.5 | 35.4 |
| F262 | -161.6 | 156.6 | F248 | -168.3 | 141.5 |
| G263 | -98.8 | 156.5 | G249 | -78.8 | 165.2 |
| K264 | -148.4 | 141.1 | K250 | -159.4 | 129.9 |
| *Lever arm* |  |  |  |  |  |
| F786 | -95.4 | 133.3 | F775 | -96.73 | 131.6 |
| F787 | -124.1 | 154.2 | F776 | -113.62 | 151.8 |
| R788 | -72.2 | 159.4 | R777 | -73.08 | 164.5 |
| A789 | -55.2 | 139.0 | T778 | -54.84 | 137.1 |
| G790 | 82.4 | -3.7 | G779 | 64.23 | 15.6 |
| V791 | -72.2 | -41.3 | V780 | -81.43 | -54.4 |
| L792 | -72.2 | -41.2 | L781 | -58.17 | -35.6 |
| A793 | -54.7 | -36.7 | A782 | -48.66 | -63.9 |
| Q794 | -78.2 | -30.3 | H783 | -53.48 | -37.6 |
| L795 | -75.2 | -32.8 | L784 | -61.41 | -42.1 |
| E796 | -67.5 | -31.7 | E785 | -63.52 | -52.9 |
| E797 | -79.4 | -32.2 | E786 | -52.85 | -42.9 |
| E798 | -74.2 | -36.6 | E787 | -66.4 | -36.1 |

**Supplementary table 2:**
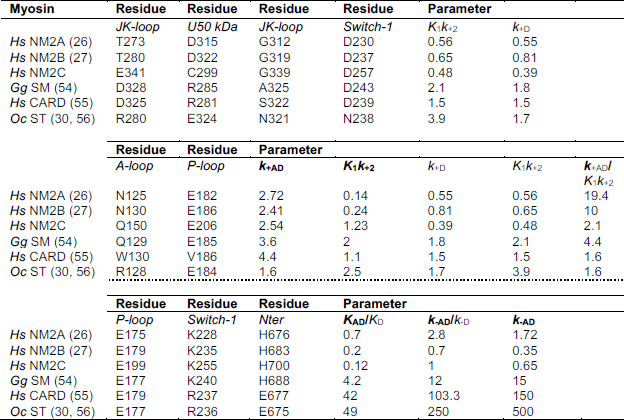
Structure function relationships in the myosin-2 motor domain. Interactions between residues in the active site involved in nucleotide binding and release kinetics based on this work and previous biochemical and structural studies on myosin motor domains (13, 16, 17, 61, 62). It is of note that myosin is a highly allosteric enzyme and nucleotide binding and release kinetic involve numerous interactions and subtle structural rearrangements of residues from different motor subdomains. Kinetic parameters from monomeric myosin motor domain constructs that are associated with a structural interaction are listed for direct comparison. An emerging trend from this analysis is that the myosin-2 kinetic cycle does not have a selectivity of ATP *versus* ADP. The presence of F-actin results in different allosteric communication pathways in myosins-2 and establishes ATP/ADP binding selectivity. Overall, nucleotide binding rates are decreased for the group of nonmuscle and smooth muscle myosins-2 compared to myosins-2 from cardiac and skeletal muscle. The lacking salt bridge interactions between JK-loop, U50 kDa and switch-1 in nonmuscle myosins-2 results in decreased second-order binding rate constants for ATP *(K*_1_*k*_+2_) and ADP *(k*_+D_) (**Figure 1 - Figure supplement 1A**). Either a salt bridge interaction or hydrophobic interactions between the A-loop and the P-loop of muscle and cardiac myosins-2 at the active site favor fast nucleotide binding kinetics and does not or only marginally discriminate between ADP and ATP. The lack of a salt bridge interactions can have different effects dependent on the coordinating residue in the A-loop: An asparagine in the A-loop of nonmuscle myosins-2A and -2B favors ADP over ATP binding to actomyosin. A glutamine in the NM2C A-loop, which has a longer side chain than asparagine, abolishes ATP/ADP sensitivity in NM2C and the closely related smooth muscle myosin-2. The number of salt bridge interactions between P-loop, switch-1, and the Nter correlates with the thermodynamic and kinetic coupling, and the actin-activated ADP release rates in all myosins-2. Abbreviations used: NM2A: human nonmuscle myosin-2A; human NM2B: nonmuscle myosin-2B (PDB entry 4PD3); NM2C: human nonmuscle myosin-2C; SM: chicken smooth muscle myosin-2 (PDB entry 1BR2); CARD: human β-cardiac myosin-2 (PDB entry 4DB1); *Oc* ST: rabbit striated muscle myosin-2 (PDB entry 1DFL).

**Supplementary table 3:**
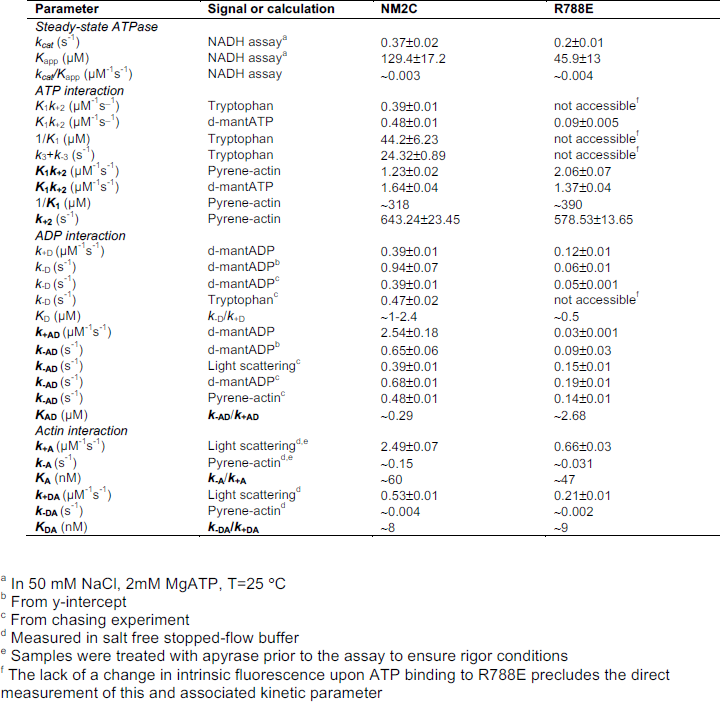
Steady-state and transient state kinetic parameters of NM2C and R788E. Numbering of the kinetic constants refers to **Figure 1 - Figure Supplement 1A**. Measurements of transient kinetic parameters that rely on a change in intrinsic tryptophan fluorescence are not experimentally accessible for R788E.

| Parameter | Signal or calculation | NM2C | R788E |
| --- | --- | --- | --- |
| *Steady-state ATPase* |  |  |  |
| $k_{cat}$ ($s^{-1}$) | NADH assay<sup>a</sup> | 0.37±0.02 | 0.2±0.01 |
| $K_{app}$ ($\mu M$) | NADH assay<sup>a</sup> | 129.4±17.2 | 45.9±13 |
| $k_{cat}/K_{app}$ ($\mu M^{-1}s^{-1}$) | NADH assay | ~0.003 | ~0.004 |
| *ATP interaction* |  |  |  |
| $K_1k_{+2}$ ($\mu M^{-1}s^{-1}$) | Tryptophan | 0.39±0.01 | not accessible<sup>f</sup> |
| $K_1k_{+2}$ ($\mu M^{-1}s^{-1}$) | d-mantATP | 0.48±0.01 | 0.09±0.005 |
| $1/K_1$ ($\mu M$) | Tryptophan | 44.2±6.23 | not accessible<sup>f</sup> |
| $k_3+k_{-3}$ ($s^{-1}$) | Tryptophan | 24.32±0.89 | not accessible<sup>f</sup> |
| $K_1k_{+2}$ ($\mu M^{-1}s^{-1}$) | Pyrene-actin | 1.23±0.02 | 2.06±0.07 |
| $K_1k_{+2}$ ($\mu M^{-1}s^{-1}$) | d-mantATP | 1.64±0.04 | 1.37±0.04 |
| $1/K_1$ ($\mu M$) | Pyrene-actin | ~318 | ~390 |
| $k_{+2}$ ($s^{-1}$) | Pyrene-actin | 643.24±23.45 | 578.53±13.65 |
| *ADP interaction* |  |  |  |
| $k_{+D}$ ($\mu M^{-1}s^{-1}$) | d-mantADP | 0.39±0.01 | 0.12±0.01 |
| $k_{-D}$ ($s^{-1}$) | d-mantADP<sup>b</sup> | 0.94±0.07 | 0.06±0.01 |
| $k_{-D}$ ($s^{-1}$) | d-mantADP<sup>c</sup> | 0.39±0.01 | 0.05±0.001 |
| $k_{-D}$ ($s^{-1}$) | Tryptophan<sup>c</sup> | 0.47±0.02 | not accessible<sup>f</sup> |
| $K_D$ ($\mu M$) | $k_{-D}/k_{+D}$ | ~1-2.4 | ~0.5 |
| $k_{+AD}$ ($\mu M^{-1}s^{-1}$) | d-mantADP | 2.54±0.18 | 0.03±0.001 |
| $k_{-AD}$ ($s^{-1}$) | d-mantADP<sup>b</sup> | 0.65±0.06 | 0.09±0.03 |
| $k_{-AD}$ ($s^{-1}$) | Light scattering<sup>c</sup> | 0.39±0.01 | 0.15±0.01 |
| $k_{-AD}$ ($s^{-1}$) | d-mantADP<sup>c</sup> | 0.68±0.01 | 0.19±0.01 |
| $k_{-AD}$ ($s^{-1}$) | Pyrene-actin<sup>c</sup> | 0.48±0.01 | 0.14±0.01 |
| $K_{AD}$ ($\mu M$) | $k_{-AD}/k_{+AD}$ | ~0.29 | ~2.68 |
| *Actin interaction* |  |  |  |
| $k_{+A}$ ($\mu M^{-1}s^{-1}$) | Light scattering<sup>d,e</sup> | 2.49±0.07 | 0.66±0.03 |
| $k_{-A}$ ($s^{-1}$) | Pyrene-actin<sup>d,e</sup> | ~0.15 | ~0.031 |
| $K_A$ (nM) | $k_{-A}/k_{+A}$ | ~60 | ~47 |
| $k_{+DA}$ ($\mu M^{-1}s^{-1}$) | Light scattering<sup>d</sup> | 0.53±0.01 | 0.21±0.01 |
| $k_{-DA}$ ($s^{-1}$) | Pyrene-actin<sup>d</sup> | ~0.004 | ~0.002 |
| $K_{DA}$ (nM) | $k_{-DA}/k_{+DA}$ | ~8 | ~9 |

<sup>a</sup> In 50 mM NaCl, 2mM MgATP, T=25 °C
<sup>b</sup> From y-intercept
<sup>c</sup> From chasing experiment
<sup>d</sup> Measured in salt free stopped-flow buffer
<sup>e</sup> Samples were treated with apyrase prior to the assay to ensure rigor conditions
<sup>f</sup> The lack of a change in intrinsic fluorescence upon ATP binding to R788E precludes the direct measurement of this and associated kinetic parameter

## References

1. Heissler SM, Sellers JR. Kinetic Adaptations of Myosins for Their Diverse Cellular Functions. Traffic. 2016;17(8):839–59.

2. Sellers JR. Myosins: a diverse superfamily. Biochim Biophys Acta. 2000;1496(1):3–22.

3. Heissler SM, Manstein DJ. Nonmuscle myosin-2: mix and match. Cell Mol Life Sci. 2013;70(1):1–21.

4. Heissler SM, Manstein DJ. Comparative kinetic and functional characterization of the motor domains of human nonmuscle myosin-2C isoforms. J Biol Chem. 2011.

5. Bloemink MJ, Geeves MA. Shaking the myosin family tree: biochemical kinetics defines four types of myosin motor. Semin Cell Dev Biol. 2011;22(9):961–7.

6. Takaoka M, Saito H, Takenaka K, Miki Y, Nakanishi A. BRCA2 phosphorylated by PLK1 moves to the midbody to regulate cytokinesis mediated by nonmuscle myosin IIC. Cancer Res. 2014;74(5):1518–28.

7. Wylie SR, Chantler PD. Myosin IIC: a third molecular motor driving neuronal dynamics. Mol Biol Cell. 2008;19(9):3956–68.

8. Ebrahim S, Fujita T, Millis BA, Kozin E, Ma X, Kawamoto S, Baird MA, Davidson M, Yonemura S, Hisa Y, Conti MA, Adelstein RS, Sakaguchi H, Kachar B. NMII forms a contractile transcellular sarcomeric network to regulate apical cell junctions and tissue geometry. Curr Biol. 2013;23(8):731–6.

9. von der Ecken J, Heissler SM, Pathan-Chhatbar S, Manstein DJ, Raunser S. Cryo-EM structure of a human cytoplasmic actomyosin complex at near-atomic resolution. Nature. 2016;534(7609):724–8.

10. Rayment I, Rypniewski WR, Schmidt-Base K, Smith R, Tomchick DR, Benning MM, Winkelmann DA, Wesenberg G, Holden HM. Three-dimensional structure of myosin subfragment-1: a molecular motor. Science. 1993;261(5117):50–8.

11. Furch M, Fujita-Becker S, Geeves MA, Holmes KC, Manstein DJ. Role of the salt-bridge between switch-1 and switch-2 of Dictyostelium myosin. J Mol Biol. 1999;290(3):797–809.

12. Reubold TF, Eschenburg S, Becker A, Kull FJ, Manstein DJ. A structural model for actin-induced nucleotide release in myosin. Nat Struct Biol. 2003;10(10):826–30.

13. Miller BM, Bloemink MJ, Nyitrai M, Bernstein SI, Geeves MA. A variable domain near the ATP-binding site in Drosophila muscle myosin is part of the communication pathway between the nucleotide and actin-binding sites. J Mol Biol. 2007;368(4):1051–66.

14. Van Driest SL, Jaeger MA, Ommen SR, Will ML, Gersh BJ, Tajik AJ, Ackerman MJ. Comprehensive analysis of the beta-myosin heavy chain gene in 389 unrelated patients with hypertrophic cardiomyopathy. J Am Coll Cardiol. 2004;44(3):602–10.

15. Havndrup O, Bundgaard H, Andersen PS, Allan Larsen L, Vuust J, Kjeldsen K, Christiansen M. Outcome of clinical versus genetic family screening in hypertrophic cardiomyopathy with focus on cardiac beta-myosin gene mutations. Cardiovasc Res. 2003;57(2):347–57.

16. Risal D, Gourinath S, Himmel DM, Szent-Gyorgyi AG, Cohen C. Myosin subfragment 1 structures reveal a partially bound nucleotide and a complex salt bridge that helps couple nucleotide and actin binding. Proceedings of the National Academy of Sciences of the United States of America. 2004;101(24):8930–5.

17. Swank DM, Braddock J, Brown W, Lesage H, Bernstein SI, Maughan DW. An alternative domain near the ATP binding pocket of Drosophila myosin affects muscle fiber kinetics. Biophys J. 2006;90(7):2427–35.

18. Sasaki N, Ohkura R, Sutoh K. Dictyostelium myosin II mutations that uncouple the converter swing and ATP hydrolysis cycle. Biochemistry. 2003;42(1):90–5.

19. Ramanath S, Wang Q, Bernstein SI, Swank DM. Disrupting the myosin converter-relay interface impairs Drosophila indirect flight muscle performance. Biophys J. 2011;101(5):1114–22.

20. Brenner B, Seebohm B, Tripathi S, Montag J, Kraft T. Familial hypertrophic cardiomyopathy: functional variance among individual cardiomyocytes as a trigger of FHC-phenotype development. Front Physiol. 2014;5:392.

21. Llinas P, Isabet T, Song L, Ropars V, Zong B, Benisty H, Sirigu S, Morris C, Kikuti C, Safer D, Sweeney HL, Houdusse A. How actin initiates the motor activity of Myosin. Dev Cell. 2015;33(4):401–12.

22. Preller M, Manstein DJ. Myosin structure, allostery, and mechano-chemistry. Structure. 2013;21(11):1911–22.

23. Bloemink MJ, Melkani GC, Bernstein SI, Geeves MA. The Relay/Converter Interface Influences Hydrolysis of ATP by Skeletal Muscle Myosin II. J Biol Chem. 2016;291(4):1763–73.

24. Kronert WA, Melkani GC, Melkani A, Bernstein SI. A Failure to Communicate: Myosin residues involved in hypertrophic cardiomyopathy affect inter-domain interaction. J Biol Chem. 2015;290(49):29270–80.

25. Kronert WA, Melkani GC, Melkani A, Bernstein SI. Mapping interactions between myosin relay and converter domains that power muscle function. J Biol Chem. 2014;289(18):12779–90.

26. Kovacs M, Wang F, Hu A, Zhang Y, Sellers JR. Functional divergence of human cytoplasmic myosin II: kinetic characterization of the non-muscle IIA isoform. J Biol Chem. 2003;278(40):38132–40.

27. Wang F, Kovacs M, Hu A, Limouze J, Harvey EV, Sellers JR. Kinetic mechanism of non-muscle myosin IIB: functional adaptations for tension generation and maintenance. J Biol Chem. 2003;278(30):27439–48.

28. Marston SB, Taylor EW. Comparison of the myosin and actomyosin ATPase mechanisms of the four types of vertebrate muscles. J Mol Biol. 1980;139(4):573–600.

29. Woodward SK, Geeves MA, Manstein DJ. Kinetic characterization of the catalytic domain of Dictyostelium discoideum myosin. Biochemistry. 1995;34(49):16056–64.

30. Ritchie MD, Geeves MA, Woodward SK, Manstein DJ. Kinetic characterization of a cytoplasmic myosin motor domain expressed in *Dictyostelium discoideum*. Proc Natl Acad Sci USA. 1993;90(18):8619–23.

31. Malnasi-Csizmadia A, Kovacs M, Woolley RJ, Botchway SW, Bagshaw CR. The dynamics of the relay loop tryptophan residue in the Dictyostelium myosin motor domain and the origin of spectroscopic signals. J Biol Chem. 2001;276(22):19483–90.

32. Friedman AL, Geeves MA, Manstein DJ, Spudich JA. Kinetic characterization of myosin head fragments with long-lived myosin. ATP states. Biochemistry. 1998;37(27):9679–87.

33. Ruppel KM, Spudich JA. Structure-function studies of the myosin motor domain: importance of the 50-kDa cleft. Mol Biol Cell. 1996;7(7):1123–36.

34. Donaudy F, Snoeckx R, Pfister M, Zenner HP, Blin N, Di Stazio M, Ferrara A, Lanzara C, Ficarella R, Declau F, Pusch CM, Nurnberg P, Melchionda S, Zelante L, Ballana E, Estivill X, Van Camp G, Gasparini P, Savoia A. Nonmuscle myosin heavy-chain gene MYH14 is expressed in cochlea and mutated in patients affected by autosomal dominant hearing impairment (DFNA4). Am J Hum Genet. 2004;74(4):770–6.

35. Fu X, Zhang L, Jin Y, Sun X, Zhang A, Wen Z, Zhou Y, Xia M, Gao J. Loss of Myh14 Increases Susceptibility to Noise-Induced Hearing Loss in CBA/CaJ Mice. Neural Plast. 2016;2016:6720420.

36. Kim BJ, Kim AR, Han JH, Lee C, Oh DY, Choi BY. Discovery of MYH14 as an important and unique deafness gene causing prelingually severe autosomal dominant non-syndromic hearing loss. J Gene Med. 2017.

37. Billington N, Wang A, Mao J, Adelstein RS, Sellers JR. Characterization of three full-length human nonmuscle myosin II paralogs. The Journal of biological chemistry. 2013.

38. Hundt N, Steffen W, Pathan-Chhatbar S, Taft MH, Manstein DJ. Load-dependent modulation of non-muscle myosin-2A function by tropomyosin 4.2. Sci Rep. 2016;6:20554.

39. Kovacs M, Thirumurugan K, Knight PJ, Sellers JR. Load-dependent mechanism of nonmuscle myosin 2. Proc Natl Acad Sci U S A. 2007;104(24):9994–9.

40. Burgess S, Walker M, Wang F, Sellers JR, White HD, Knight PJ, Trinick J. The prepower stroke conformation of myosin V. J Cell Biol. 2002;159(6):983–91.

41. Heissler SM, Chinthalapudi K, Sellers JR. Kinetic characterization of the sole nonmuscle myosin-2 from the model organism Drosophila melanogaster. FASEB J. 2015;29(4):1456–66.

42. Blessing RH. An empirical correction for absorption anisotropy. Acta Crystallogr A. 1995;51(Pt 1):33–8.

43. McCoy AJ. Solving structures of protein complexes by molecular replacement with Phaser. Acta Crystallogr D Biol Crystallogr. 2007;63(Pt 1):32–41.

44. Emsley P, Cowtan K. Coot: model-building tools for molecular graphics. Acta Crystallogr D Biol Crystallogr. 2004;60(Pt 12 Pt 1):2126–32.

45. Bricogne G, Blanc E, Brandl M, Flensburg C, Keller P, Paciorek W, Roversi P, Sharff A, Smart OS, Vonrhein C, Womack TO. autoBUSTER. Cambridge: Global Phasing Ltd. 2011.

46. Davis IW, Murray LW, Richardson JS, Richardson DC. MOLPROBITY: structure validation and all-atom contact analysis for nucleic acids and their complexes. Nucleic Acids Res. 2004;32(Web Server issue):W615–9.

47. Malnasi-Csizmadia A, Toth J, Pearson DS, Hetenyi C, Nyitray L, Geeves MA, Bagshaw CR, Kovacs M. Selective perturbation of the myosin recovery stroke by point mutations at the base of the lever arm affects ATP hydrolysis and phosphate release. J Biol Chem. 2007;282(24):17658–64.

48. MacKerell AD, Bashford D, Bellott M, Dunbrack RL, Evanseck JD, Field MJ, Fischer S, Gao J, Guo H, Ha S, Joseph-McCarthy D, Kuchnir L, Kuczera K, Lau FT, Mattos C, Michnick S, Ngo T, Nguyen DT, Prodhom B, Reiher WE, Roux B, Schlenkrich M, Smith JC, Stote R, Straub J, Watanabe M, Wiorkiewicz-Kuczera J, Yin D, Karplus M. All-atom empirical potential for molecular modeling and dynamics studies of proteins. J Phys Chem B. 1998;102(18):3586–616.

49. Phillips JC, Braun R, Wang W, Gumbart J, Tajkhorshid E, Villa E, Chipot C, Skeel RD, Kale L, Schulten K. Scalable molecular dynamics with NAMD. J Comput Chem. 2005;26(16):1781–802.

50. Jorgensen WL, Chandrasekhar J, Madura JD, Impey RW, Klein ML. Comparison of Simple Potential Functions for Simulating Liquid Water. J Chem Phys. 1983;79(2):926–35.

51. Darden TA, Pedersen LG. Molecular modeling: an experimental tool. Environ Health Perspect. 1993;101(5):410–2.

52. Foth BJ, Goedecke MC, Soldati D. New insights into myosin evolution and classification. Proc Natl Acad Sci U S A. 2006;103(10):3681–6.

53. Sellers JR. Myosins: a diverse superfamily. Biochim Biophys Acta. 2000;1496(1):3–22.

54. Cremo CR, Geeves MA. Interaction of actin and ADP with the head domain of smooth muscle myosin: implications for strain-dependent ADP release in smooth muscle. Biochemistry. 1998;37(7):1969–78.

55. Deacon JC, Bloemink MJ, Rezavandi H, Geeves MA, Leinwand LA. Identification of functional differences between recombinant human alpha and beta cardiac myosin motors. Cellular and molecular life sciences: CMLS. 2012.

56. Kurzawa-Goertz SE, Perreault-Micale CL, Trybus KM, Szent-Gyorgyi AG, Geeves MA. Loop I can modulate ADP affinity, ATPase activity, and motility of different scallop myosins. Transient kinetic analysis of S1 isoforms. Biochemistry. 1998;37(20):7517–25.

57. Wagner PD. Formation and characterization of myosin hybrids containing essential light chains and heavy chains from different muscle myosins. J Biol Chem. 1981;256(5):2493–8.

58. Harris DE, Warshaw DM. Smooth and skeletal muscle myosin both exhibit low duty cycles at zero load in vitro. J Biol Chem. 1993;268(20): 14764–8.

59. Kliche W, Fujita-Becker S, Kollmar M, Manstein DJ, Kull FJ. Structure of a genetically engineered molecular motor. EMBO J. 2001;20(1-2):40–6.

60. Munnich S, Pathan-Chhatbar S, Manstein DJ. Crystal structure of the rigor-like human non-muscle myosin-2 motor domain. FEBS Lett. 2014;588(24):4754–60.

61. Grammer JC, Kuwayama H, Yount RG. Photoaffinity labeling of skeletal myosin with 2-azidoadenosine triphosphate. Biochemistry. 1993;32(22):5725–32.

62. Szilagyi L, Balint M, Sreter FA, Gergely J. Photoaffinity labelling with an ATP analog of the N-terminal peptide of myosin. Biochem Biophys Res Commun. 1979;87(3):936–45.

